# Dendritic cell actin dynamics controls T cell priming efficiency at the immunological synapse

**DOI:** 10.1101/2020.06.13.150045

**Authors:** Alexander Leithner, Lukas M. Altenburger, Robert Hauschild, Frank Assen, Klemens Rottner, Theresia E.B. Stradal, Jens V. Stein, Michael Sixt

## Abstract

Dendritic cells (DCs) are crucial for the priming of naïve T cells and the initiation of adaptive immunity. Priming is initiated at a heterologous cell-cell contact, the immunological synapse (IS). While it is established that actin dynamics regulates signalling at the T cell side of the contact, little is known about the cytoskeletal contribution on the DC side. We show that that the DC cytoskeleton is decisive for the formation of a multifocal synaptic structure, which correlates with T cell priming efficiency. We demonstrate that DC actin appears in transient foci at the IS and that these foci are dynamized by the WAVE complex. Absence of WAVE in DCs leads to stabilized contacts with T cells, caused by an increase in ICAM1-integrin mediated cell-cell adhesions. This results in a lower number of activated and proliferating T cells. Our results reveal an important role of DC actin in the regulation of synaptic contacts with crucial relevance for full T cell expansion.

## Introduction

Dendritic cells (DCs) are the most potent antigen presenting cells (APCs) with the unique ability to prime naïve T cells *in vivo* (Banchereau and Steinman, 1998). T cell activation requires the formation of a transient cell-cell contact between T cell and APC. Within this immunological synapse (IS) (Monks et al., 1998; Dustin et al., 1998), the T cell receptor (TCR) together with co-stimulatory molecules and adhesion receptors engage their ligands on the APC surface to trigger downstream signaling, resulting in T cell activation and proliferation. The T cell face of the IS has been extensively studied in experimental setups where the APC is replaced by supported lipid bilayers (SLBs), which allow for imaging with high spatio-temporal resolution (Grakoui et al., 1999). On SLBs the T cell IS is organized in the classic monofocal configuration, comprising three concentric domains: the central-, peripheral-, and distal supramolecular activation cluster (c-, p-, and dSMAC). While the cSMAC was initially thought to be the site of TCR signaling, it turned out to be the center of TCR recycling (Varma et al., 2006; Das et al., 2004). Instead, signaling occurs *en route*, when TCR microclusters (MCs) that arise in the dSMAC travel with a centripetal actin flow towards the cSMAC (Babich et al., 2012; Yi et al., 2012). The pSMAC forms an adhesive ring that stabilizes the IS, mediated by the integrin lymphocyte function-associated antigen 1 (LFA-1) that binds its ligand intercellular adhesion molecule 1 (ICAM1) on the APC surface.

While the IS between B cells, which have weak priming capacity (Gunzer et al., 2004), and T cells resembles the aforementioned monofocal configuration found on SLBs, the IS between DCs and T cells is more complex and, despite its crucial physiological capacity, poorly understood (Fisher et al., 2008). T cell-DC ISs were described as multifocal, exhibiting multiple local ensembles of TCR, co-receptors and adhesion molecules (Brossard et al., 2005; Tseng et al., 2008). In line with the idea that this multifocal structure determines the superior T cell priming capacity of DCs, engagement of the co-stimulatory molecule CD28 in multiple peripheral, instead of one central, clusters correlates with stronger T cell activation (Shen et al., 2008).

The molecular driver of differential IS patterning is currently unknown. In the monofocal IS, formed between T and B cells and natural killer cells- or cytotoxic T cells and target cells, the cytoskeleton seems largely cleared from the presynaptic face (Friedl et al., 2005). Therefore, SLBs where ligands freely float in the bilayer, appear to be a valid model for such “passive” APCs. This is different in DCs, where the F-actin cytoskeleton polarizes towards the IS (Al-Alwan and Rowden, 2001; Benvenuti et al., 2004), supporting the idea that DCs actively pattern the IS via their cytoskeleton in order to support optimal T cell priming (Dustin et al., 2006a; Comrie and Burkhardt, 2016). As the presynaptic face of the IS has received little attention, direct evidence for this idea is missing. DC F-actin might form a stable scaffold but could potentially also have a more dynamic role in structuring the IS.

Here we use a multimodal imaging approach, in combination with functional *in vitro* and *in vivo* assays, to investigate the role of presynaptic actin dynamics in DCs, and its impact on the priming of naïve T cells.

## Results & Discussion

To explore how the structure of the T cell IS is influenced by the DC actin cytoskeleton, we decided to depolymerize F-actin in DCs and to characterize synapses formed with freshly isolated CD4+ T cells from OT-II transgenic mice whose T cells carry a TCR specific for an Ovalbumin-derived peptide (Barnden et al., 1998). In order to selectively target actin in DCs and leave T cells unperturbed, we pre-treated DCs with the drug mycalolide B (mycB). MycB causes complete and lasting actin depolymerization by severing F-actin filaments and irreversible sequestration of G-actin (Hori et al., 1993; Saito et al., 1994; Vaahtomeri et al., 2017). In contrast to other drugs like cytochalasin D that has been used in earlier studies (Al-Alwan and Rowden, 2001), mycB leads to covalent modification and therefore cannot be washed out. MycB treated DCs lost their F-actin rich veils, adopted a spherical shape and stained negative for phalloidine (Fig.1A & C). Pre-treated DCs underwent a slight shift towards lower surface levels of the DC markers Cd11c and MHCII (Fig.1B) with no apparent effect on cell viability (Supplementary Figure 1A & B). Synapses between mycB-treated or control DCs and T cells were fixed, stained for F-actin and imaged via confocal microscopy. Around 50% of synapses between F-actin free DCs and untreated T cells exhibited a monofocal structure with a pronounced ring of F-actin surrounding a central region devoid of F-actin. In contrast, all synapses in control samples showed a multifocal structure (Fig.1C & D). These results suggest that the DC actin cytoskeleton prevents formation of a monofocal IS, potentially by providing barriers that block centripetal flow of T cell actin (Dustin et al., 2006b). However, we also note that upon DC F-actin depolymerization only about 50% of synapses become clearly monofocal, indicating that also other, F-actin independent, DC properties may contribute to IS patterning.

**Figure 1.**
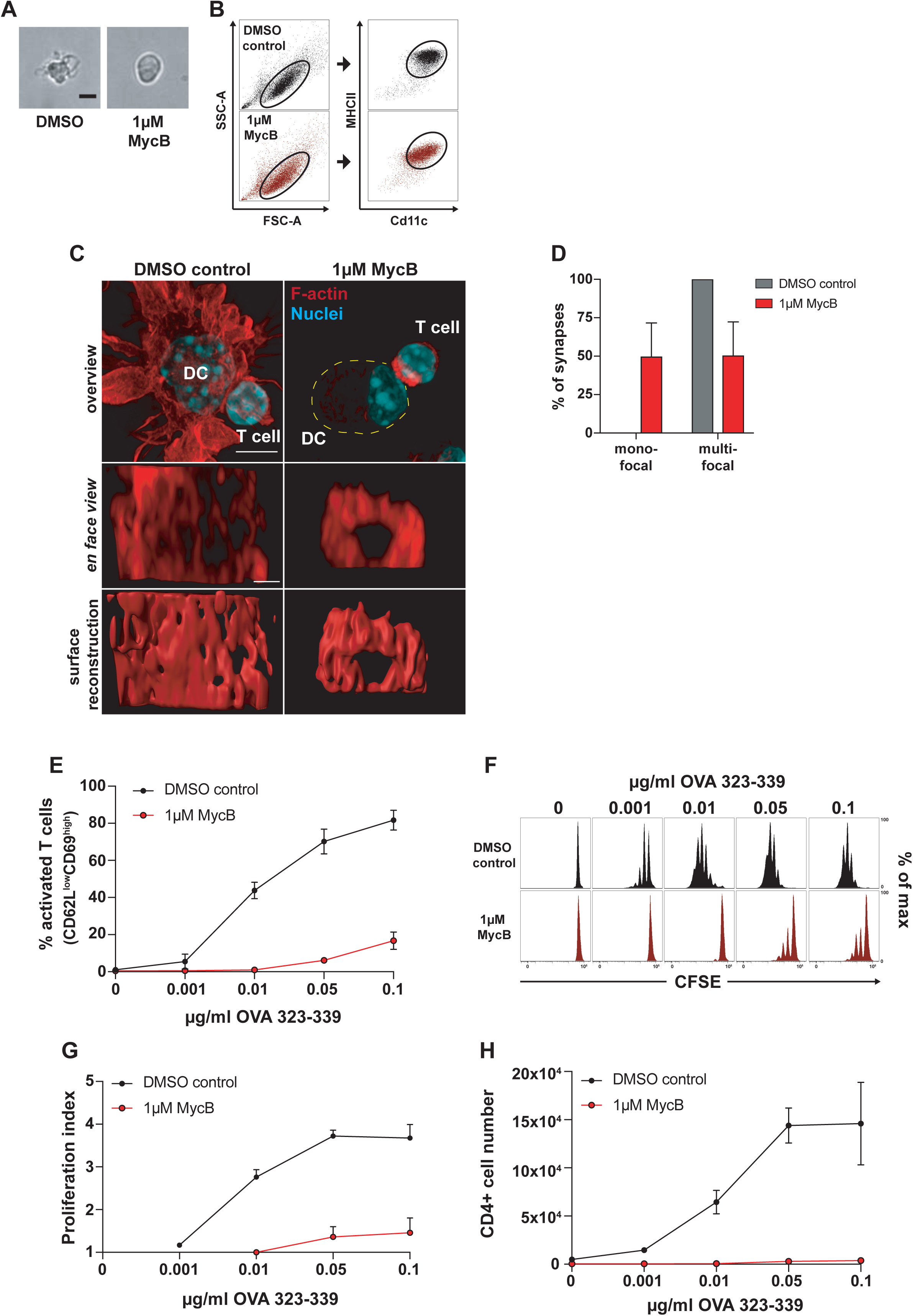
DC F-actin depolymerisation affects immune synapse structure and T cell priming efficiency. (A) Bright field images of DCs treated with 1µM mycB (right) or DMSO (left), scale bar = 10µm. (B) Flow cytometry profiles of DMSO and 1µM mycB-treated DCs. Cd11c/MHCII plots are pre-gated on FSC-A/SSC-A population defined by black oval on the left, representative example of three biological replicates. (C) Immunofluorescence images of synapses formed between DMSO or 1µM mycB-treated DCs and T cells; upper panel: overview pictures, yellow dotted line outlines DC, scale bar = 5µm; middle panel: *en face* view on the synaptic interface, scale bar = 1µm; lower panel: surface reconstruction of the synaptic interface. (D) Percentages of mono- and multifocal synapses formed between T cells and 1µM mycB-treated DCs, n∼20 cell duplets for each condition, 3 biological replicates, mean ± SD. (E) Percentages of activated T cells assessed by CD62L/CD69 surface expression after 16h of co-culture with DMSO or 1µM mycB-treated DCs at indicated OVA_323-339_ peptide concentrations, 3 biological replicates, mean ± SD. (F) CFSE dilution profile of T cells after 96h of co-culture with DMSO or 1µM mycB-treated DCs at indicated OVA_323-339_ peptide concentrations, representative example of 3 biological replicates. (G) Proliferation indices of CFSE-labelled T cells after 96h of co-culture with DMSO or 1µM mycB-treated DCs at indicated OVA_323-339_ peptide concentrations, 3 biological replicates, mean ± SD. (H) Absolute T cell numbers after 96h of co-incubation with DMSO or 1µM mycB-treated DCs at indicated OVA_323-339_ peptide concentrations, 3 biological replicates, ± SD.

Next, we determined how actin depolymerization in DCs affects T cell priming. The ability of mycB pre-treated DCs to activate T cells was dramatically decreased as measured by surface levels of early activation markers CD62L and CD69 (Fig. 1E). Impaired activation resulted in strongly reduced T cell proliferation with few T cells entering division even at very high peptide concentrations (Fig.1F-H). Taken together, these results suggest that the DC actin cytoskeleton shapes the structure of the synapse, which is intricately linked to the strength of T cell activation. Importantly, these results are in line with observations in SLB systems where addition of artificial barriers that prevent monfocal synapse formation enhance TCR signaling (Mossman et al., 2005).

Next, we were interested *how* DC actin at the IS might support T cell activation. Besides the fact that DC actin accumulates at the IS, little is known about its structure, dynamics and molecular regulation. This is mainly due to the difficulty to image dynamic cell-cell contacts *en-face* and the complication that in stained samples actin from the pre- and postsynaptic face blurs. To circumvent this problem, we developed a setup where antigen-loaded, Lifeact-eGFP expressing DCs and fluorophore-labelled T cells are confined between a cover-glass and a layer of polydimethylsiloxane (PDMS) (le Berre et al., 2014). In this narrow space, DCs and T cells frequently position on top of each other and form synapses within the horizontal imaging plane (Fig.2A). Z-plane reconstruction of fast spinning disc confocal movies shows that DC actin accumulates at the cell-cell interface as suggested previously from side-views of fixed samples (Fig.2B, right). However, visualization of the synaptic plane revealed that DC actin dynamically appears in small foci and larger patches, which are separated by actin-depleted regions. Notably, there was no indication of any centripetal actin flow (Fig.2B, C & F).

**Figure 2.**
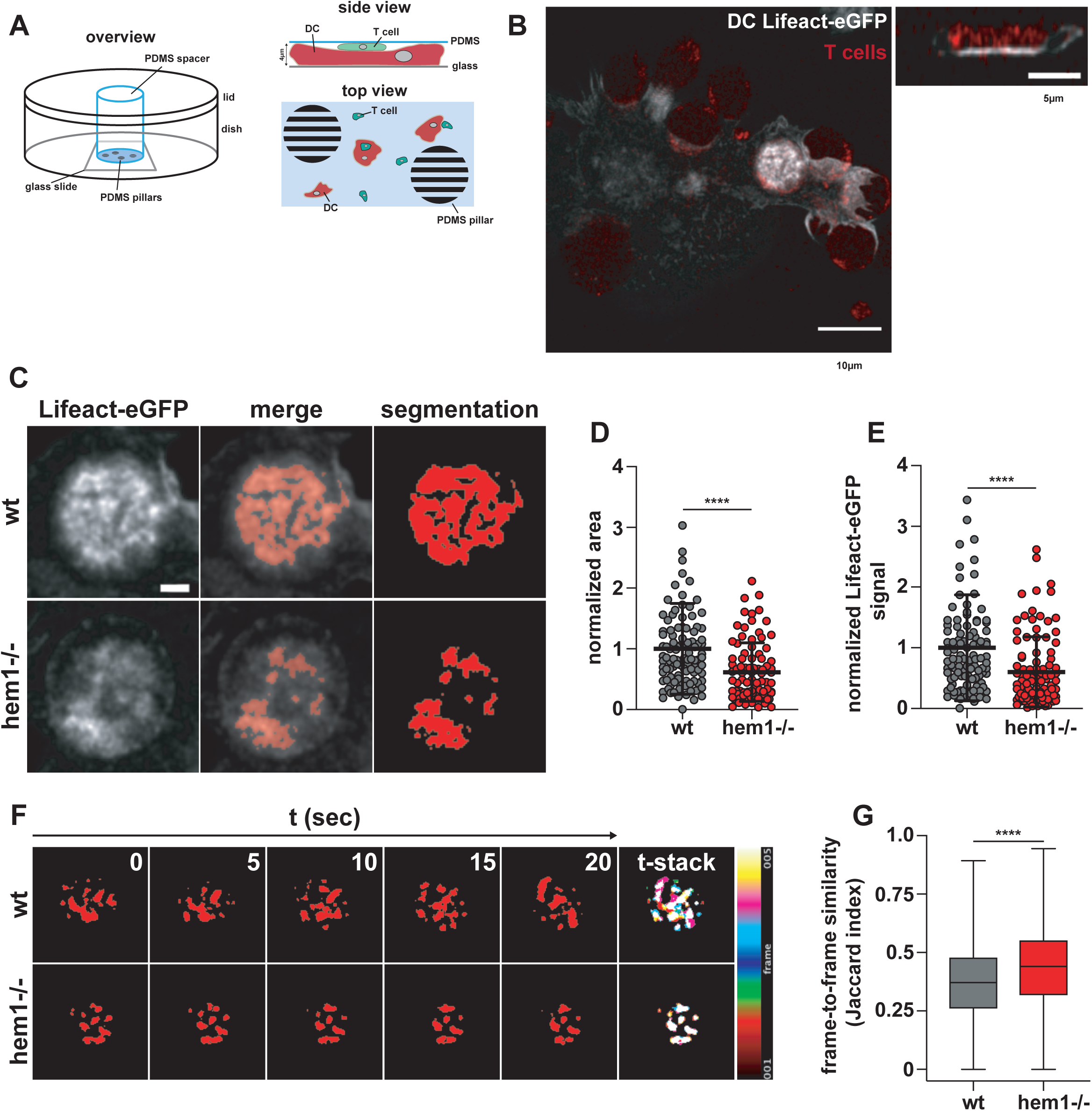
Dynamic F-actin foci at the DC immune synapse. (A) Schematic overview of PDMS confiner setup. (B) *Left*: spinning disc snapshot of Lifeact-eGFP-expressing wt DC interacting with TAMRA-stained T cells, scale bar = 10µm, *right*: z-stack of bright Lifeact-eGFP signal region on the left, scale bar =5µm. (C) Examples of wt and hem1-/- DC-T cell immune synapses and machine learning-based segmentation of synaptic DC Lifeact-eGFP signal, scale bar = 2µm. (D) Normalized area of synaptic DC Lifeact-eGFP signal, n∼100 synapses each, Mann-Whitney test p<0.0001, mean ± SD, 3 biological replicates. (E) Normalized synaptic DC Lifeact-eGFP signal, n∼100 synapses each, Mann-Whitney test p<0.0001, mean ± SD, 3 biological replicates. (F) *Left*: Time-lapse series of synaptic DC Lifeact-eGFP signal, *right*: stack through time-lapse series. (G) Frame-to-frame similarity of synaptic DC Lifeact-eGFP signal, n∼6000 frame comparisons each, Mann-Whitney test p<0.0001, mean ± min/max, 2 biological replicates.

We next addressed the molecular regulation of synaptic actin in DCs. WAVE, together with WASP, are the main nucleation promoting factors (NPFs) for the Arp2/3 complex that nucleates branched actin networks at the plasma membrane (Takenawa and Suetsugu, 2007; Alekhina et al., 2017). At the T cell face of the IS, WAVE and WASP have critical and divergent functions. WASP polymerizes F-actin foci that are associated with TCR MCs that recruit PLCγ1, leading to calcium influx and NFAT signaling. In contrast, WAVE creates and reorganizes the synaptic actin background that mediates integrin-dependent adhesion and PLCγ1-independent signaling (Nolz et al., 2006; Kumari et al., 2015). In DCs, WASP has been shown to be important for T cell-DC contact formation and duration and IS stability (Bouma et al., 2011; Pulecio et al., 2008; Malinova et al., 2016). As the role of DC WAVE at the IS is unknown, we generated WAVE-deficient DCs from bone marrow of mice lacking *hem1*, an essential subunit of the pentameric WAVE complex specific for the hematopoietic system (Park et al., 2008). Hem1-/- DCs contain 50% less F-actin, while surface levels of MHCII and co-stimulatory ligands are equal to control cells (Leithner et al., 2016). We used machine learning-based image segmentation (Berg et al., 2019) to perform a quantitative and unbiased comparison of actin dynamics in the synapses of wt vs hem1-/- DCs. Actin occupied a significantly reduced area-fraction of the contact surface in hem1-/- compared to control DCs (Fig.2C, D & E). This effect was not due to altered levels of the Lifeact-eGFP probe that is expressed at similar levels in wt and hem1-/- DCs (Supplementary Fig.1C). When comparing time-lapse movies of synaptic wt and hem1-/- DC Lifeact-eGFP, we found that actin foci of wt DCs frequently appear and disappear throughout the whole synaptic interface. In contrast, by comparing the frame-to-frame similarity of the Lifeact-eGFP signal, we found that hem1-/- DC actin foci display significantly reduced lateral dynamics (Fig.2F & G and Supplementary movie 1).

Next, we asked how WAVE deficiency affects the DC’s T cell priming capacity. We found that compared to control cells hem1-/- DCs were only able to activate a small fraction of T cells at early time-points, as measured by modulation of early activation markers (Fig.3A & B). This deficiency in T cell priming was prominent even under the highest peptide concentrations, accompanied by reduced IL-2 levels in the supernatant (Fig.3C) and resulted in significantly fewer T cells after 4 days of DC T cell co-culture (Fig. 3D & E). However, in dye dilution assays, which measure proliferation at the single cell level, we could not detect any differences in the proliferative indices of T cells stimulated by wt or hem1-/- DCs (Fig. 3F). This indicates that, although hem1-/- DCs activate fewer T cells, the ones that are activated proliferate normally. As these data indicated that changes in contact strength or frequency might reduce the capacity of DCs to prime multiple T cells, we analyzed the relative contact area of T cells with DCs in fixed duplets. Morphometry revealed that hem1-/- DCs and T cells form significantly larger contacts compared to wt DCs and that hem1-/- DCs often wrap around T cells covering a substantial part of their surface (Fig.4A & B). These findings led us to speculate that the reduced presynaptic actin dynamics in WAVE-deficient DCs translates into an overall reduction in dynamism of cell-cell contacts. To test this idea, we seeded wt or hem1-/- DCs and T cells on glass slides and quantified their interactions using live cell microscopy. Contact times between OVA peptide loaded hem1-/- DCs and T cells were substantially increased compared to the wt control, while in the absence of peptide, interaction times were equally low (Fig.4C and Supplementary movie 2).

**Figure 3.**
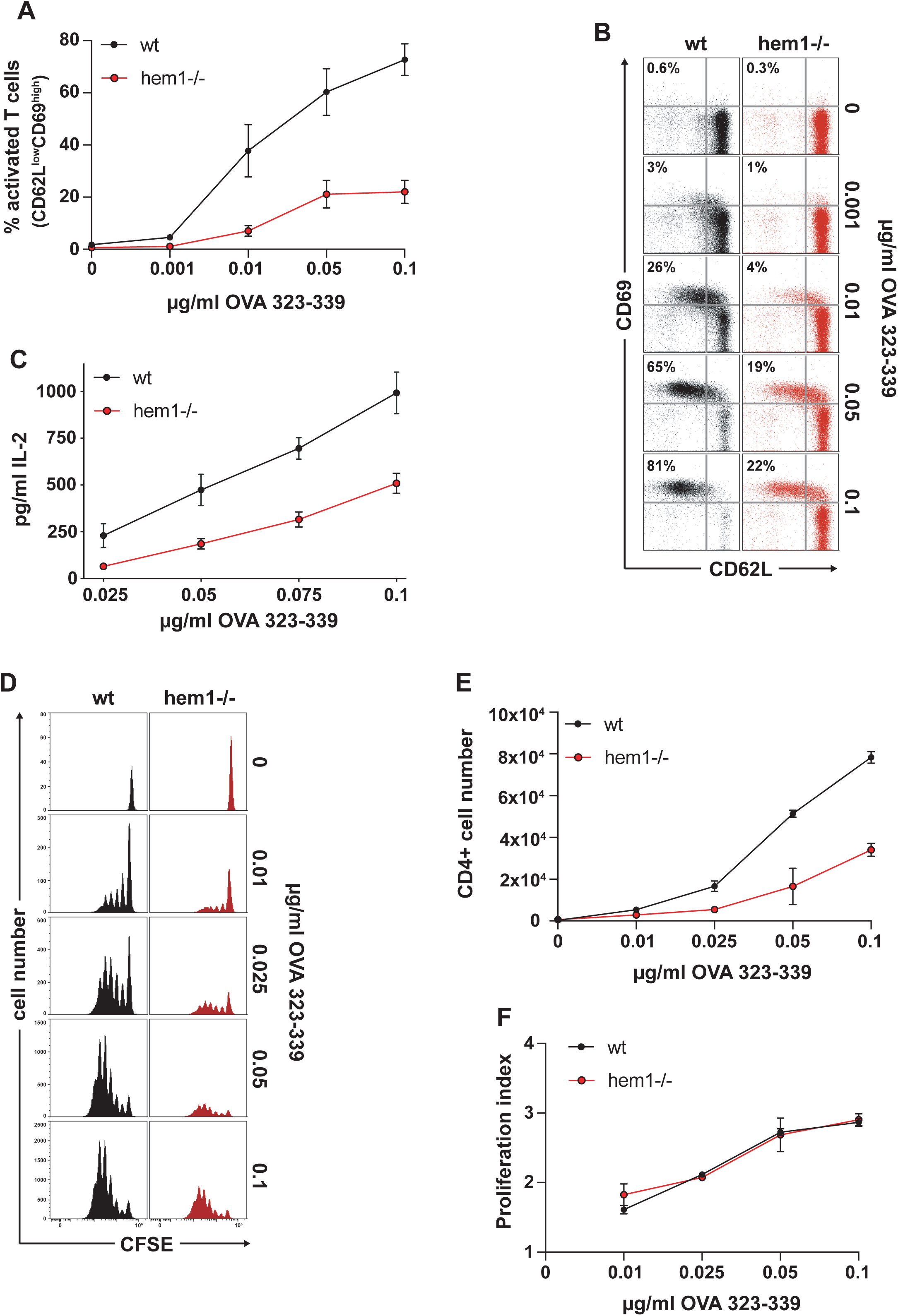
Hem1-/- DCs are impaired in T cell activation. (A) Percentage of activated T cells assessed by CD62L/CD69 surface expression at indicated OVA_323-339_ peptide concentrations, 3 biological replicates, mean ± SD. (B) Exemplary CD62L/CD69 flow cytometry profile of T cells after 16h of co-culture with wt or hem1-/- DCs. (C) IL-2 ELISA after 16h of T cell-wt or hem1-/- DC co-culture at indicated OVA_323-339_ peptide concentrations, 3 biological replicates, mean ± SD. (D) CFSE dilution profile of T cells after 96h of co-culture with wt or hem1-/- DCs at indicated OVA_323-339_ peptide concentrations, representative example of 3 biological replicates. (E) Absolute T cell numbers after 96h of co-incubation with wt or hem1-/- DCs at indicated OVA_323-339_ peptide concentrations, 3 biological replicates, ± SD. (F) Proliferation indices of CFSE-labelled T cells after 96h of co-culture with wt or hem1-/- DCs at indicated OVA_323-339_ peptide concentrations, 3 biological replicates, mean ± SD.

**Figure 4.**
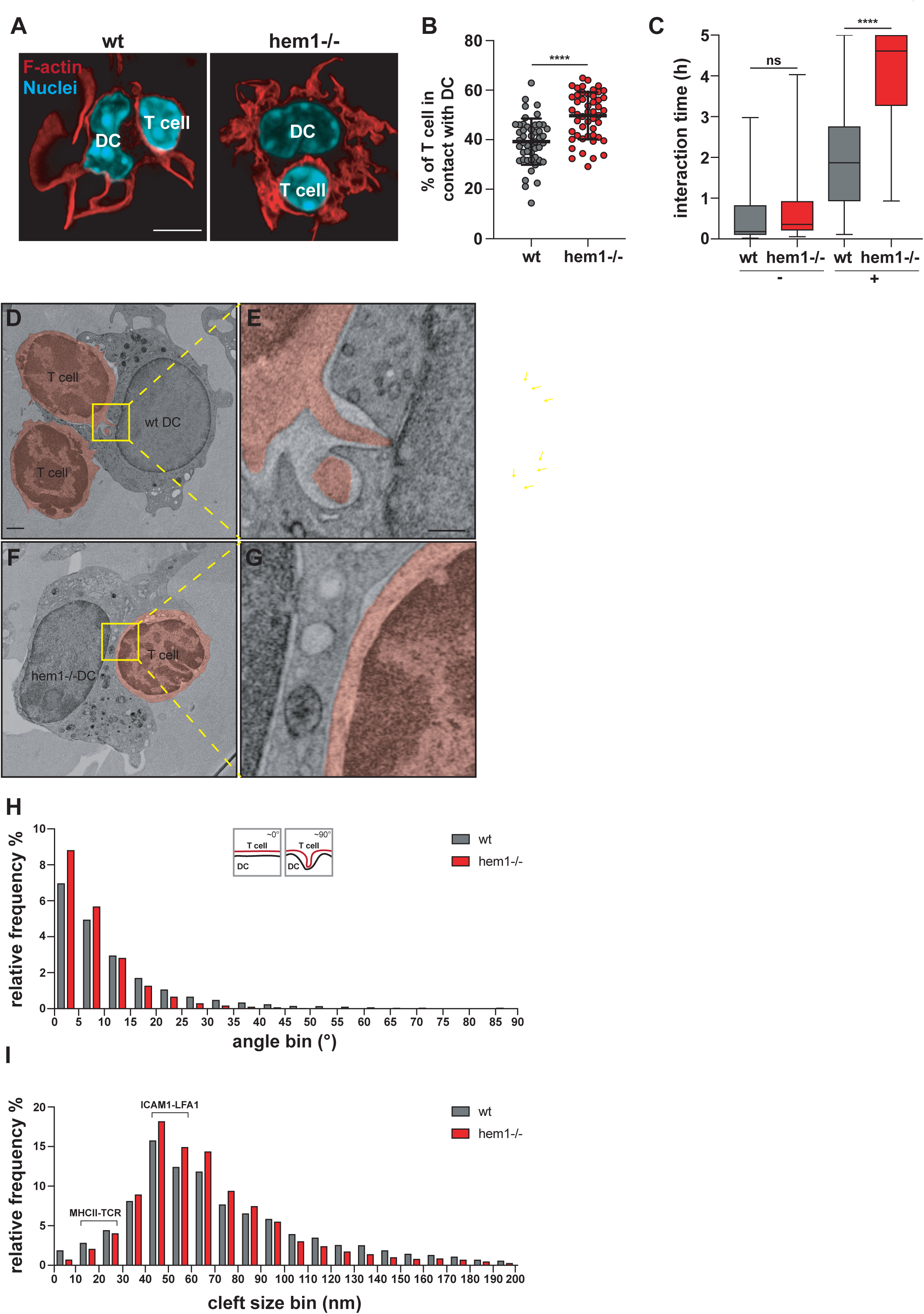
Hem1-/- DCs alter synapse structure and dynamics. (A) Immunofluorescence images of synapses formed between wt or hem1-/- DCs and T cells stained with phalloidin and DAPI, scale bar = 5µm. (B) Percentages of T cell surface area in contact with DC, ∼50 cells each, t-test p<0.0001, mean ± SD, 3 biological replicates. (C) Interaction times of wt or hem1-/- DCs with T cells in the absence (-) or presence (+) of OVA_323-339_ peptide, n∼50 contacts each for (+) or n∼ 30 contacts each for (-), Mann-Whitney test p<0.0001, mean ± min./max., 3 biological replicates. (D) Electron micrograph of wt DC-T cell synapse, T cells are coloured in red, yellow box and dotted lines denote region magnified in (E), (F) Electron micrograph of hem1-/- DC-T cell synapse, T cell is coloured in red, yellow box and dotted lines denote region magnified in (G), scale bars=1µm in (D) & (F), 300nm in (E) and (G). (H) Frequency histograms in percent of the angles found between DC and T cell membranes, n∼4 synapses each, 2 biological replicates. (I) Frequency histograms in percent of the cleft size found between DC and T cell membranes from (H).

To understand the mechanistic cause of these stabilized contacts we resolved the ultrastructure of the whole synaptic interface by performing electron microscopy of high pressure-frozen and serially sectioned DC-T cell duplets. The wt DC-T cell synapse is characterized by multiple finger-like protrusions emanating from T cells and protruding into the DC’s cell body. At these focal protrusions the plasma membranes of both cells come into direct contact (Fig.4D & E and Supplementary movie 3). Similar protrusions have been observed between T cells and other cell types (Ueda et al., 2011; Sage et al., 2012) and might correspond to T cell ‘microvilli’ that have been recently observed on artificial substrates (Cai et al., 2017; Jung et al., 2016). In contrast, hem1-/- DC-T cell contacts are characterized by a complete lack of T cell protrusions, giving the T cell membrane at the synapse a smooth and stretched appearance (Fig.4F & G and Supplementary movie 4). To quantify this difference, we traced the plasma membranes of DCs and T cells and determined at which angle they are positioned relative to each other. Morphometry revealed a shift towards lower angles for hem1-/- DC-T cell synapses demonstrating parallel alignment of both membranes. Wt DC-T cell synapses exhibit higher angles as membranes position orthogonal in regions of T cell protrusions (Fig. 4H).

The synaptic cleft was previously shown to be differentially spaced depending on the functional zone of the synapse. Sites of integrin-mediated adhesion are separated by 35-55nm, matching the large extracellular parts of LFA1 and ICAM1. The shorter TCR-MHC pair translates into closer contacts below 25nm (Shaw and Dustin, 1997). Thus, we determined the distances between DC and T cell membranes and found an overrepresentation of 35-55nm distances and an underrepresentation of below 25nm distances in Hem1-/- compared to control synapses (Fig.4I).

These data suggested that ICAM1-LFA1 interactions were more frequent in hem1-/- DC-T cell synapses. Integrin adhesiveness is regulated by their clustering, termed valency, and their ligand affinity (Kinashi, 2005). They engage in catch-bonds where application of pulling force on the integrin-ligand pair shifts integrins into their high affinity conformation (Kong et al., 2009; Chen et al., 2010; Zhu et al., 2008). Consequently, it has been suggested that forces are provided by the T cell F-actin cytoskeleton (Zhu et al., 2008; Schürpf and Springer, 2011) and that these forces have to be balanced by opposing retention forces. In support of this idea only immobilized, but not soluble, ICAM1 triggers conversion of LFA-1 to the high affinity conformation (Perez et al., 2003; Feigelson et al., 2010).

As surface ICAM1 levels of hem1-/- DCs were unchanged (Supplementary Figure 1D), we turned our attention to the adaptor proteins that connect ICAM1 to the underlying F-actin cytoskeleton. In DCs, surface ICAM1 was shown to be immobilized by anchorage to the F-actin cytoskeleton via the phosphorylated and thus active form of the ERM protein moesin. This enables DCs to oppose forces that are exerted on LFA-1 through the T cell F-actin cytoskeleton in a cellular ‘tug of war’, making them well suited to facilitate transition of LFA-1 into its high affinity conformation and thus support full T cell activation (Comrie et al., 2015). We found that pERM levels are elevated in hem1-/- compared to wt DCs while total ERM levels were unchanged (Fig.5A & B). Elevated pERM levels immobilize ICAM1, potentially causing an increase in LFA-1 valency at the hem1-/- DC-T cell synapse. This might stabilize the synapse, counter its resolution and thereby lead to increased interaction times.

**Figure 5.**
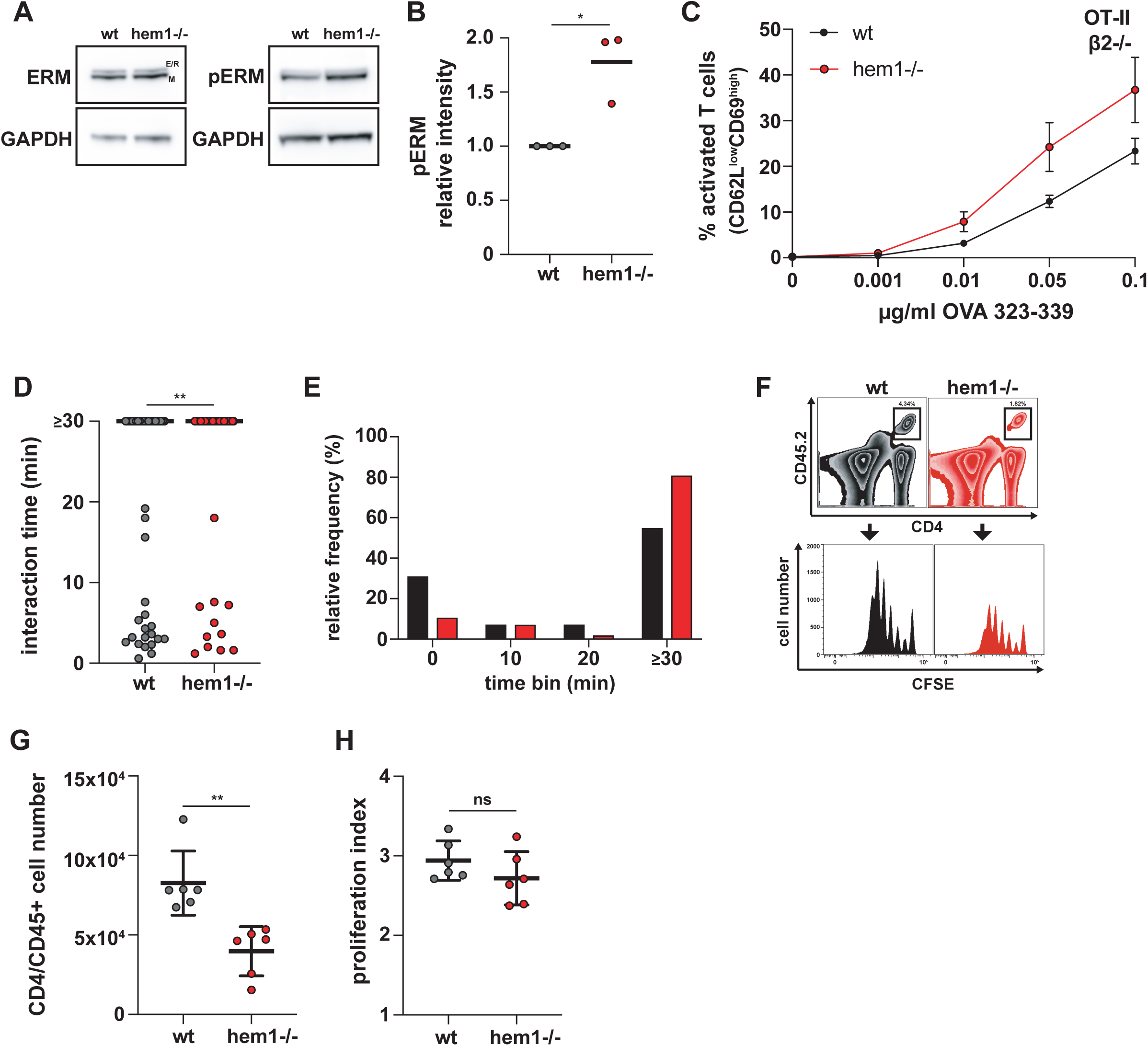
Hem1-/- DCs have T cell priming defects in vivo, mediated by the pERM-ICAM1-LFA1 axis. (A) Western blots for ERM, pERM and GAPDH in wt and hem1-/- DCs, representative example of 3 biological replicates. (B) Relative intensity of pERM signal in wt and hem1-/- DCs, t-test p=0.0158, 3 biological replicates. (C) Percentages of activated beta 2 integrin-deficient T cells assessed by CD62L/CD69 surface expression at indicated OVA_323-339_ peptide concentrations, 3 biological replicates, mean ± SD. (D) *In vivo* interaction times of wt or hem1-/- DCs and T cells, n∼45 cell-cell interactions each, Mann-Whitney test p=0.0064, bars indicate median, 2 biological replicates. (E) Frequency histograms in percent of DC-T cell interaction times from (D). (F) Gating strategy to determine the CFSE proliferation profiles of CD45.2/CD4 T cells in CD45.1 mice after wt or hem1-/- DC T cell priming. (G) Absolute CD45.2/CD4 T cell numbers in the popliteal lymph nodes of CD45.1 mice 72h after DC injections, n=6 lymph nodes each, Mann-Whitney test p=0.0022, 2 biological replicates. (H) Proliferation indices of CFSE-labelled T cells from (G), t-test not significant.

If the T cell priming deficiency of hem1-/- DCs is caused by an increase of adhesiveness, disruption of the ICAM1-LFA-1 axis should rescue this effect. To test this idea, we co-cultured wt and hem1-/- DCs together with naïve T cells of integrin beta 2 deficient, OTII transgenic mice and monitored T cell activation. Removal of ICAM1-LFA-1 interactions led to an overall reduction of T cell activation (compare Fig.3A and Fig.5C). Notably, under these conditions the hem1-/- DCs were no more inferior in their priming capacity. In contrast, they even triggered slightly enhanced activation of integrin-deficient T cells when compared to their wt counterparts. This suggests that stabilized cell-cell adhesion via the ICAM1-LFA-1 axis is the main determinant for the T cell priming deficiency of hem1-/- DCs.

While the importance of the ICAM1-LFA-1 axis for *in vitro* T cell priming has been established (Abraham et al., 1999), its *in vivo* role remains controversial. Deletion of LFA-1 or talin in T cells leads to strong *in* vivo T cell proliferation defects (Kandula and Abraham, 2004; Wernimont et al., 2011). However, these results have to be interpreted with care due to the severe homing defects of these T cells. In contrast, deletion of ICAM1 on lymph node-resident (Feigelson et al., 2018) and migratory DCs (Kozlovski et al., 2019) appears to have no effect on initial T cell activation and proliferation but is crucial for the development of immunological memory (Scholer et al., 2008). To address these points, we performed hock co-injections of differently labelled and peptide-loaded wt and hem1-/- DCs, followed by intravenously injected labelled OT-II T cells. This procedure mimics an infectious setting and causes DCs to home into the lymph node via the lymphatic route, while T cells enter the node via the blood circulation. We then performed two-photon intravital microscopy of the popliteal lymph node and quantified cellular interactions. Contact times between hem1-/- DCs and T cells were significantly increased compared to wt, reminiscent of the *in vitro* situation (Fig.5D & E and Supplementary movies 5&6). Finally, to determine if this change in contact times has an effect on T cell proliferation we performed *in vivo* CFSE dilution assays. While activated T cells underwent the same number of divisions, significantly less T cells accumulated with hem1-/- DCs, again mirroring our *in vitro* findings (Fig.5F-H).

How does increased adhesion at the hem1-/- DC-T cell IS lead to a reduction in T cell priming? Formation of a stable IS has initially been considered a requirement for full T cell activation (Dustin et al., 1997). However, later studies have shown that, specifically when T cells are activated by DCs, phases of short, sequential contacts and a dynamic synaptic interface (kinapse) are the prevalent pattern (Bousso, 2008; Gunzer et al., 2000; Dustin, 2008). We suggest that balancing cell-cell adhesion and de-adhesion, regulated by WAVE and WASP on the DC-side, is an important factor in achieving full T cell expansion. DC-T cell contacts have to be long enough for T cells to become fully activated but interaction time has to be regulated in a way that allows to contact as many T cells as possible. These results are in line with previous studies where DC adhesion to T cells was artificially increased by activation of normally inactive DC MAC-1 or LFA-1, leading to impaired T cell priming (Varga et al., 2007; Balkow et al., 2010). Alternatively, increased ICAM1-LFA-1 interaction might lead to suboptimal spacing of the DC and T cell membranes. This potentially counteracts segregation of the TCR-MHC pair that is required for optimal T cell activation (Comrie and Burkhardt, 2016).

## Acknowledgments

We thank the Electron microscopy facility, the Bioimaging facility, the Pre-clinical facility and the Life Science facility of IST Austria for excellent support. This work was supported by the European Research Council (ERC CoG 724373) and a grant from the Austrian Science Fund (FWF P29911) to M.S.. K.R. and T.EB.S. are funded by the German Research Council (DFG) and the Helmholtz Society.

## Materials & Methods

### Cell culture

R10 medium, consisting of RPMI 1640 supplemented with 10% fetal calf serum (FCS), 2mM L-glutamine, 100U/ml penicillin, 100µg/ml streptomycin and 50µM 2-mercaptoethanol (all Thermo Fisher) was used as basic cell culture medium.

*Dendritic cells* were differentiated from bone marrow of male or female 6-12-week-old wt or hem1-/- C57BL/6J mice according to established protocols (Lutz et al., 1999). F-actin reporter mice were obtained by breeding wt or hem1+/- mice with Lifeact-eGFP animals (Riedl et al., 2010), followed by back-crosses to hem1+/- mice. 2×10^6^ wt or 1.25×10^6^ hem1-/- bone marrow cells were seeded in 9ml R10, supplied with 1ml in-house-generated granulocyte-macrophage colony stimulating factor (GM-CSF) into 94mm petri dishes (Greiner, 632180). At day 3, 8ml R10, supplied with 2ml GM-CSF was added to the dishes. At day 6, 10ml of medium were removed and replaced with 8ml R10 and 2ml GM-CSF. On day 8 or 9, non-adherent cells from two dishes were collected and cryo-preserved in 1ml 90% DMSO and 10% FCS. For maturation, two vials were thawed and cells were seeded overnight onto 150mm cell culture dishes (VWR, 734-2322) in 18ml R10 and 2ml GM-CSF, supplied with 200ng/ml lipopolysaccharide (LPS) from *E.coli* 0127:B8 (L4516, Merck). Only cells from the supernatant were used in experiments. For some experiments, cells were treated with 1µM mycB (enzo lifesciences, BML-T123) for 15min and then washed three times with R10 medium.

*T cells* were isolated from spleens and lymph nodes of C57BL/6J OTII transgenic mice (Barnden et al., 1998) by negative CD4+ selection (Stemcell, 19852). LFA-1-deficient T cells were obtained by breeding CD18-/- (Wilson et al., 1993) to OT-II mice. For these mice, negative selection for naïve CD4+ was performed (Stemcell, 19765). T cells were directly used for experiments after isolation.

### DC marker staining

0.5×10^6^ DCs/tube were spun down (300g, 5min.), washed once- and then re-suspended in FACS buffer (1xPBS, 5mM EDTA, 1% BSA). Cells were first stained with Fc-block (1:100, clone 93) and then with a mixture of anti-mouse Cd11c-APC (1:300, clone N418) and anti-mouse MHCII-eFluor450 (1:800, clone M5/114.15.2, all ebioscience). All stainings were performed at 4°C for 15min. Cells were washed once with 1ml FACS buffer, re-suspended and analyzed on a FACS Canto II machine (BD).

### Imaging of live and fixed DC-T cell duplets

Glass bottom dishes (35mm, 14mm glass diameter, glass thickness 0, Mattek) were plasma cleaned (pdc-002 plasma cleaner, Harrick) and coated with 1x poly-L-lysine (P8920, Merck) in H_2_O for 10min. Dishes were washed twice with H_2_O and then dried for at least 4h at room temperature. DCs were pre-loaded with 0.1µg/ml OVA_323-339_ in R10 (vac-isq, invivogen) for 2h at 37°C/5% CO_2_. 1.5×10^5^ DCs were mixed with 3×10^5^ T cells and then pipetted onto coated dishes in a total volume of 300µl. Cells were either followed by live imaging for 5h or allowed to interact for 30min at 37°C/5% CO_2_, followed by fixation with 300µl 6% PFA in R10. Cells were then washed twice with 1xTBS and permeabilized with 0.2% TritonX-100 in 1x TBS for 10min. Samples were washed twice and then blocked with 3% BSA in 1xTBS for 1h at room temperature. F-actin was stained with phalloidin-Alexa488 (1:40, A12379, Thermofisher) for 30min, washed twice with 1xTBS and then embedded in mounting medium with DAPI (Fluoromount-G with DAPI, 00-4959-52, Thermofisher). Samples were imaged on a LSM880 with Airyscan (Zeiss).

### *In vitro* T cell activation, proliferation and IL-2 assays

The assays were carried out in 96-well, round bottom plates (TPP). 10.000 DCs and 50.000 T cells per well were mixed in 200µl R10 in the presence of different concentrations of OVA_323-339_ peptide.

To assess T cell activation, 16-18h after initiation of the co-culture, plates were spun down (350g, 10min., 4°C), and the supernatants removed and snap-frozen. Cells were stained with Fc block and then with a mixture of anti-mouse CD62L-PE (1:1500, MEL-14), anti-mouse CD69-APC-eFluor780 (1:200, H1.2F3) and anti-mouse CD4-eFluor450 (1:800, GK1.5, all ebioscience) in FACS buffer.

To measure T cell proliferation, T cells were stained with 5µM CFSE (C1157, Thermofisher) as previously described (Quah et al., 2007). 96h after initiation of the co-culture, cells were spun down as described above and stained for CD4. 7AAD was used to distinguish live from dead cells. Data were recorded with a FACS Canto II machine (BD), equipped with an automated high throughput sampler. FlowJo (www.flowjo.com) was used for data analysis.

The frozen supernatants were used for IL-2 ELISAs according to the manufacturer’s protocol (KMC0021, Thermofisher).

### *In vivo* T cell proliferation

CD45.1 ‘pepboy’ mice were injected intravenously with 1×10^6^ freshly isolated CD4/CD45.2 OTII T cells, stained with CFSE as described above. 24h later, DCs, pre-loaded with 1µg/ml OVA_323-339_ peptide were injected into the hind footpad. As hem1-/- DCs have a disadvantage in reaching the draining lymph node compared to wt DCs (Leithner et al., 2016), we injected double the amounts of hem1-/- DCs (4×10^5^), which leads to comparable cell numbers in the lymph node (data not shown). 72h after DC injection, mice were sacrificed and popliteal lymph nodes isolated. Lymph nodes were torn open and digested for 30min at 37°C in digestion buffer (DMEM + 2%FCS, 3mg/ml Collagenase IV, 40µg/ml DNAseI, 3mM CaCl_2_). Cells were filtered through a cell strainer and washed and stained in FACS buffer with anti-mouse CD4-eFluor450 (1:300, clone GK1.5, ebioscience) and anti-mouse CD45.2-APC (1:300, clone 104, ebioscience).

### Intravital microscopy

DCs were pre-loaded with 2µM OVA_323-339_ peptide for 45min. at 37°C/5% CO_2_. In the last 20min. of loading, hem1-/- DCs were labelled with 25µM CMTMR (C2927, Thermofisher), wt DCs with 20µM CMAC (C2110, Thermofisher). Cells were washed once, mixed in a 2:1 ratio (hem1-/- : wt) and 1.5×10^6^ cells in 20µl were injected into the hock. 18h post injection, 3×10^6^ GFP expressing OT-II T cells were injected intravenously in a volume of 100µl. Reactive popliteal lymph nodes were imaged by intravital two-photon microscopy in 30min. intervals, 2-8h after T cell transfer.

### Electron microscopy

Sapphire discs were coated with carbon land markings and incubated with 1x poly-L-lysine (P8920, Merck) in H_2_O over night at 4°C. On the next day, discs were placed in a 12-well dish, rinsed twice with H_2_O and dried for 4h at room temperature. wt or hem1-/- DCs, pre-loaded with 0.1μg/ml OVA_323-339_ peptide for 2h together with T cells were seeded in R10 and allowed to interact for 2h. Samples were mildly pre-fixed with 1% PFA for 10min and then immediately transferred to a Baltec HPM010 machine for high pressure freezing. Freeze substitution was carried out in an automated Leica EMAFS machine according to the following protocol: 23.5h at -82°C with acetone and 0.1% tannic acid, followed by a change to acetone, 1% osmium tetroxide and 0.2% uranylacetate and incubation for another 7h. The temperature was then raised to -60°C over a time period of 1.5h, kept constant for 3h and then raised to -30°C over the course of 2h, followed by a final rise to 0°C. Embedding was started by acetone washes and infiltration of the samples with resin/propylene oxide mixtures with increasing amounts of resin. Samples were infiltrated with pure resin overnight and then allowed to polymerize at 60°C for 2-3 days. Serial sections of 70nm were cut with an automated tape collecting ultra-microtome (ATUMtome, RMC) and placed on a waiver for semi-automated image collection with a Merlin compact VP field-emission scanning electron microscope (Zeiss). The Fiji plugin ‘TrakEM2’ (Cardona et al., 2012) was used for image alignment. To quantify the distances between DC and T cell membranes, the T cell volume was segmented manually using Microscopy Image Browser (Belevich et al., 2016). The DC membrane was drawn as a ROI. A simple Fiji script was then used to calculate the binary distance map from the T cell and to evaluate the distances at the location of the DC membrane (ROI). The angle between DC and T cell membranes was quantified with a custom script. To this end, the manually drawn ROI of the DC membrane as well as the manual segmentation of the T cell was used. Contour lines of the T cell are constructed as the normal of the gradient of the distance map of the T cell segmentation. The freehand line of the ROI that marks the DC membrane is cleaned up and a cubic smoothing spline is fitted to it. For the entire length of the membrane, the angle between the contour line and the membrane was calculated at steps of 100nm.

### Western blots

1×10^6^ wt and hem1-/- DCs were spun down and lysed in 50µl 1x lysis buffer (1ml 10x RIPA buffer - NEB, 9ml H_2_O, 1 tablet PhosStop and 1 tablet protease inhibitor mini - both Merck). Lysates were spun at full speed for 10min at 4°C and supernatants transferred to new tubes. Equal volumes of 2x loading buffer (4x LDS buffer, 10x sample reducing agent) were added. Two lanes were run for each wt or hem1-/- sample. Per lane, 15µl were denatured at 99°C for 5min. Samples were loaded on a 10% Tris-glycine gel (Thermofisher) and run with 180V. After protein transfer, membranes were cut in a way to allow separate incubation with anti-ERM (1:1000, #3142, cell signaling) anti-pERM (1:1000, #3141, cell signaling) and anti-GAPDH (1:10.000, ab125247, abcam) antibodies. Membranes were blocked with 5% BSA in 1xTBS-T for 1h at room temperature, followed by incubation with the respective antibodies overnight at 4°C. Membranes were washed three times for 10min with 1xTBS-T and then incubated with HRP-coupled anti-rabbit secondary antibodies (170-6515, Biorad) for 1h at room temperature. Membranes were washed again for three times with 1xTBS-T, incubated with SuperSignal West Femto Maximum Sensitivity Substrate (Thermofisher) and imaged on an Amersham imager 600 (GE healtcare).

### Manufacturing of and imaging in PDMS confiner

In brief, the imaging chamber consists of two glass surfaces that are spaced by polydimethylsiloxane (PDMS) micropillars. The two surfaces are pressed together by a PDMS piston that is glued onto the lid of a glass bottom dish. The photomask design for the PDMS micropillars was drawn with Coreldraw X8 (Corel) and printed on an emulsion film transparency with a resolution of 8mm (JD Photo Data and Photo tools). The mold was produced by photolithography on a silicon wafer. In brief, the wafer was coated with SU8-GM1050 (Gersteltec) at 2.120rpm for 40s to achieve a height of 4µm. The wafer was soft-baked for 1min at 120°C and for 5min at 95°C and then exposed to UV light at 1mJ/cm^2^ for 10min using a beam-expanded 365nm UV LED (M365L2-C1-UV, Thorlabs). After UV exposure, the wafer was post-baked for 1min at 65°C and 5min at 95°C. The wafer was developed in SU8 developer for 17s and then silanized with trichloro (1H, 1H, 2H, 2H-perfluorooctyl) silane in a vacuum desiccator for 1h. Micropillars were then produced by mixing silicone elastomer and curing reagent (PDMS Sylgard 184 Elastomere Kit, Dow Corning) in a ratio of 7:1. The mixture was then degassed, using a planetary centrifugal mixer (ARE250, Thinky), and carefully poured on the wafer. Round cover glasses (#1, 12mm diameter, Mentzel, Thermofisher) were plasma activated for 2min. (Plasma Cleaner, Harrick Plasma) and placed on the wafer with the activated surface facing the elastomere/curing agent mixture. The wafer was cured on a heating plate for 15min at 95°C and the micropillar-coated cover glasses were carefully removed with a sharp razor blade and isopropanol. To produce the PDMS piston, silicone elastomer and curing reagent were mixed in a 30:1 ratio, degassed as described before and poured in an aluminum mold with the needed dimensions. The PDMS pistons were cured for 6h at 80°C, removed with isopropanol and then glued onto the middle of the lid of a 60×15mm non-pyrogenic polystyrene tissue culture dish (Falcon), using aquarium sealant (Marina). A hole with a diameter of 17mm was drilled in the center bottom of a 60×15mm tissue culture dish (Falcon) and a glass slide (#2, 22×22mm, Mentzel, Thermofisher) was glued onto the bottom of the hole with aquarium sealant.

To assemble the device, a micropillar-bearing cover glass was mounted on the PDMS piston with the micropillars facing upward. DCs and T cells were mixed in a volume of 5µl that was then carefully pipetted on the micropillar. The PDMS piston with the micropillars and cell mixture was then pressed onto the glass slide in the tissue culture dish that was then sealed with strong tape. Dishes were incubated for 1h at 37°C/5% CO_2_ before imaging was performed on an inverted confocal microscope (Zeiss) equipped with a spinning-disc system (iXon897, Andor), a Plan-Apochromat 100×/1.4 oil objective (Zeiss) and 488nm and 561nm lasers.

Still images of the synapse were obtained by z-stacking in 27, ∼300nm intervals. The Huygens Deconvolution Software package (https://svi.nl/Huygens-Deconvolution) was used for deconvolution according to the manufacturer’s manual. The synaptic plane was then identified as the one with the highest Lifeact-eGFP signal and together with one plane above and below was used to obtain maximum intensity projections that were used for further analysis. Live imaging was performed by recording three, ∼133nm-spaced z-planes at the synaptic interface in a 5sec interval for 3min. Maximum intensity projections were used for further analysis. To quantify F-actin and its dynamics, we first segmented the actin-rich area in the synapse using ilastik (Berg et al., 2019). The segmentation was manually corrected and used to extract area and intensity measures from the raw data. All intensity measures were background corrected. Area and intensity measures were normalized to the respective mean of wt DCs. To quantify F-actin dynamics, single pixels where automatically removed. As a measure for how much the actin segmentation changes from frame to frame, we calculated the Jaccard similarity coefficient. It is defined by the overlap of the segmentation area of the current frame with the segmentation of the previous frame divided by the area of the union of both. Hence, a completely static synapse would lead to a Jaccard coefficient of 1 and a lower number indicates a more dynamic synapse.

### Statistics and data analysis

All data were analyzed using Graph Pad Prism 8.

## Figure Legends

**Supplementary Figure 1.**
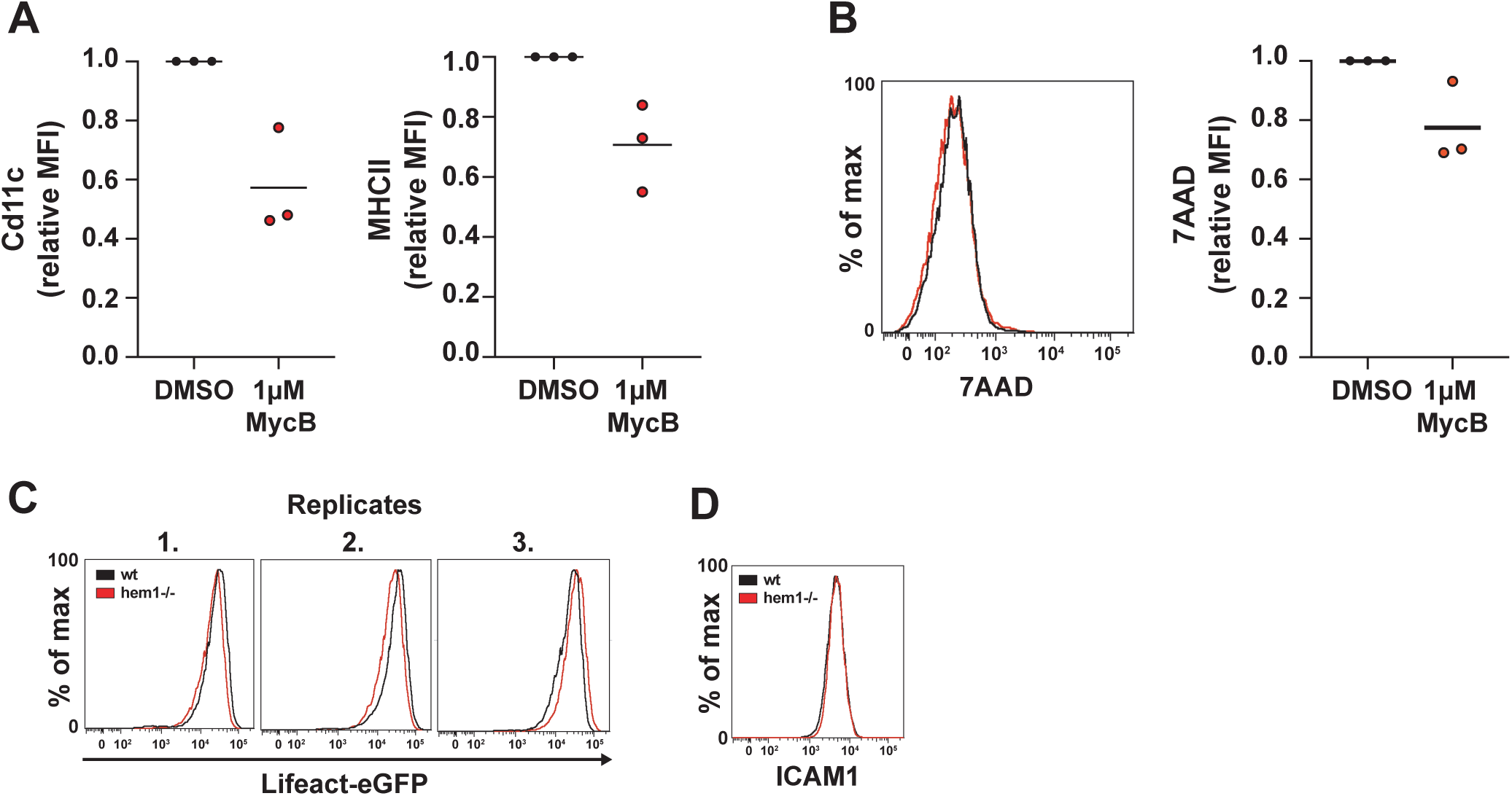
(A) Flow cytometry experiment displaying relative mean fluorescence intensity for Cd11c (left) and MHCII (right) of DMSO or 1µM mycB-treated DCs, 3 biological replicates (B) 7AAD life/dead stain flow cytometry histogram (left) and relative mean fluorescence intensity (right) of DMSO or 1µM mycB-treated DCs, 3 biological replicates. (C) Flow cytometry histograms of 3 biological replicates for Lifeact-eGFP expressing wt and hem1-/- DCs (D) Flow cytometry histogram for ICAM1 in wt and hem1-/- DCs.

## Movie Legends

**Supplementary movie 1.** Lifeact-eGFP signal of OVA peptide-loaded wt or hem1-/- DCs forming immune synapses with OT-II T cells in confined setup; *Left*: raw data, *right*: segmentation; frame interval: 5sec, time stamp: min:sec, scale bar: 5µm.

**Supplementary movie 2.** Interactions of OVA peptide-loaded wt or hem1-/- DCs with OT-II T cells on poly-l-lysine coated glass; frame interval: 30sec, time stamp: h:min, scale bar: 10µm.

**Supplementary movie 3.** Z-stack through high pressure-frozen and serially sectioned synapse formed between OVA peptide-loaded wt DC and OT-II T cell (red); z-interval: 30nm, scale bar: 1µm.

**Supplementary movie 4.** Z-stack through high pressure-frozen and serially sectioned synapse formed between OVA peptide-loaded hem1-/- DC and OT-II T cell (red); z-interval: 30nm, scale bar: 1µm.

**Supplementary movie 5.** Intravital microscopy of interactions between OVA peptide-loaded wt DCs (blue) and OT-II T cells (green); frame interval: 20sec, time stamp: min:sec, scale bar: 5µm.

**Supplementary movie 6.** Intravital microscopy of interactions between OVA peptide-loaded hem1-/- DCs (red) and OT-II T cells (green); frame interval: 20sec, time stamp: min:sec, scale bar: 5µm.

